# Minimally invasive brain injections for viral-mediated transgenesis: New tools for behavioral genetics in sticklebacks

**DOI:** 10.1101/2020.03.02.973594

**Authors:** Noelle James, Alison Bell

## Abstract

Behavioral genetics in non-model organisms is currently gated by technological limitations. However, with the growing availability of genome editing and functional genomic tools, complex behavioral traits such as social behavior can now be explored in diverse organisms. Here we present a minimally invasive neurosurgical procedure for a classic behavioral, ecological and evolutionary system: threespine stickleback (*Gasterosteus aculeatus*). This method of direct brain injection enables viral-mediated transgenesis and pharmaceutical delivery which bypasses the blood-brain barrier. This method is flexible, fast, and amenable to statistically powerful within-subject experimental designs, making it well-suited for use in genetically diverse animals such as those collected from natural populations.

Viral-mediated transgenesis in the brain allows for a direct examination of the genetic mechanisms underlying behavior in wild-caught animals from natural populations. Using this method, we were able to detect changes in aggression from the knockdown of either of two different genes, arginine vasopressin (*AVP*) and monoamine oxidase (*MAOA*), in outbred animals in less than one month. In addition, we demonstrate that widely available mammalian plasmids work with this method, lowering the barrier of entry to the technique.

## Introduction

Complex behaviors have been repeatedly shown to be heritable (reviewed in Dochtermann et al., 2019), yet establishing a causal relationship between genes and social behavior remains challenging. Partially, this difficulty arises from limitations of the primarily correlative methods for examining the interplay between genes and behavior (reviewed in Charney 2017). While DNA sequence changes were historically thought to rely on changes in protein efficacy, they can occur in promoter or enhancer regions resulting in changes in expression (Bossdorf et al., 2007; Xie et al., 2019). Changes in gene expression are frequently lumped together under the category of epigenetics which rely on changes outside the DNA sequence such as methylation or histone modifications.

DNA sequence differences giving rise to different phenotypes can be identified with techniques such as quantitative trait locus (QTL) mapping and genome-wide linkage/association studies (GWAS). RNA-Seq, currently one of the most popular methods for identifying candidate genes, detects both sequence-based and epigenetic effects, including alternative splicing and post-transcriptional modifications. These techniques yield candidate genes from regions where variation in genetic and epigenetic markers is significantly associated with the phenotypic variation. To fully characterize how a gene contributes to behavior, it is necessary to consider not just sequence differences, but also regulatory and epigenetic influences. Therefore, to demonstrate and fully characterize a causal relationship between a gene and behavior, it is crucial to have a method for manipulating gene expression at a specific time and location (London, 2020).

It is important to select the right tools for the system as each method has its own assumptions and required extant capabilities within the system (Bengston et al., 2018). Pharmacological manipulation can mimic the results of altered gene expression (Stefanska and MacEwan, 2015). However, pharmaceutical manipulation is more useful in examining the signaling mechanisms at the protein level, while directly altering gene expression would be preferable for establishing a causal relationship between genes and behavior. Viral-mediated transgenesis is a method to increase a gene’s expression in a specific location or time (Ingusci et al., 2019; Simonato et al., 2000; Zou et al., 2014). This approach has already proved essential in the functional testing of genes related to behavior (Gallant and O’Connell, 2020; London, 2020; Simonato et al., 2000) as well as in the dissection of neural circuits (Luo et al., 2008).

Viral-mediated transgenesis can be used to increase or decrease gene expression levels, using either a direct gene payload or a CRISPR cassette, respectively (Braasch et al., 2014; Ingusci et al., 2019). This method of transgenesis is fast and flexible and amenable to experimental designs in which the same individual animals are measured before and after gene expression is altered, allowing us to show a causal relationship. By using a repeated measure within-subject design, each animal acts as its own control, which statistically eliminates variation between individuals. This is a major benefit when dealing with high inter-individual variation, such as when studying behavior or outbred animals collected from natural populations.

To enable direct manipulation of candidate genes and thereby examine how they contribute to behavior, we developed a minimally invasive neurosurgical method for direct brain injection in the threespine stickleback (*Gasterosteus aculeatus*). This procedure can deliver either pharmacological agents to test signaling pathways or transgenic elements to directly alter gene expression into the stickleback brain. Threespine stickleback have long been a foundational organism in ethology (reviewed in Huntingford & Ruiz-Gomez 2009). Already one of the best-studied animals for behavior, sticklebacks have a growing molecular toolkit including a fully sequenced genome (Jones et al., 2012).

Thus, sticklebacks are now are gaining popularity in other fields including evolution, physiology, comparative genomics, and neuroscience (Fang et al., 2018; Norton and Gutiérrez, 2019). They have been used in comparative cross-taxa studies looking for a conservation in the molecular underpinnings of social behavior with both emerging and classic model systems (Rittschof et al., 2014; Saul et al., 2019), as well as in the evolution of behavior (Di Poi et al., 2016; Dingemanse et al., 2012; Fang et al., 2018; Huntingford and Ruiz-Gomez, 2009). Indeed, there are already hundreds of previously identified candidate genes for social behavior waiting to be characterized (Bell et al., 2016; Bukhari et al., 2017; Greenwood and Peichel, 2015; Greenwood et al., 2013; Laine et al., 2012; Mommer and Bell, 2014; Sanogo et al., 2011). Sticklebacks enjoy well-established behavioral assays (Rowland, 1982; van Iersel, 1953) that are amenable to automation (Ardekani et al., 2013; Norton and Gutiérrez, 2019).

There is a dearth of information on surgical methodology in small (3-4 cm) fish, despite their popularity as behavioral model systems, including zebrafish (*Danio rerio*) (Maruska et al., 2019), medaka (*Oryzias latipes*) (Kirchmaier et al., 2015), and stickleback (Huntingford and Ruiz-Gomez, 2009). Anesthetization remains one of the most challenging aspects of surgery because proper anesthetizations timing must be determined on an individual basis. In modifying a zebrafish viral-mediated transgenesis procedure (Zou et al., 2014), we first needed to refine the anesthesia process (Neiffer and Stamper, 2009; Sladky and Clarke, 2016) to enable a longer and more precise surgery. Importantly, we introduce the use of an oral cannula providing oxygenation and maintenance anesthetic throughout the longer (10 minutes out-of-water) procedure. To improve the precision of brain injections, we designed and built a custom surgical rig as there are no commercially-available, water compatible, tiny stereotaxis tables. To maximize animal welfare, we additionally needed to identify clear warning signs of failure to recuperate by establishing a normal recovery pattern in stickleback similar to the work in koi by Harms et al., (2005). With this equipment and the ability to provide early intervention, survival rates rose to 90%.

To facilitate future behavioral work with this stereotactic brain surgery for stickleback, we tested methods to minimize post-surgical downtime such as supplemental oxygenation and verified prompt behavioral recovery via a simulated territorial intrusion immediately after physiological recovery. As the first test of this method in this species, we chose to focus on territorial aggression for three reasons: 1) it is a well-established, easy to score behavioral assay (Bell, 2005; Norton and Gutiérrez, 2019; Rowland and Bolyard, 2000; van Iersel, 1953; Wootton, 1971), 2) aggression is important for fitness (Anholt and Mackay, 2012; Huntingford and Turner, 1987), and 3) there are good candidate genes for aggression based on studies in other vertebrates (Sanogo et al., 2012; Saul et al., 2019; Takahashi and Miczek, 2014).

Here we employ this neurosurgical method to test the function of two conserved candidate genes related to aggression in stickleback: arginine-vasopressin and monoamine oxidase. Using viral-mediated transgenesis via direct brain injections, we tested if behavior level changes were detectable due to brain-wide overexpression of either *AVP* or *MAOA* in wild-caught male sticklebacks. To increase the accessibility of viral-mediated transgenesis in sticklebacks, a system with roots in ethology rather than genetics, we demonstrate the use of the technique for behavioral genetics by using widely available mammalian plasmids, rather than synthesizing or cloning out the stickleback gene sequences. To confirm the efficacy of pharmaceutical manipulations via brain injections, we compared brain and systemic injection of vasotocin.

Arginine-vasopressin (*AVP*) and its nonmammalian homolog arginine-vasotocin (*AVT*) are highly conserved (Moore, 1992) and pleiotropic (Balment et al., 2006). Vasopressin and vasotocin are distinguished by only a single amino acid change between mammals (human) and teleosts (sticklebacks), and their respective V1a receptors have similar specificity, signaling mechanisms, and amino acid sequences (Goodson and Bass, 2001). Both vasopressin and vasotocin were found to have similar physiological effects in rats (Feuerstein et al., 1984). Additionally, vasotocin signaling has been shown to influence aggression in various contexts in both fish and mammals (reviewed in Goodson, 2013) and has been characterized throughout the social decision making network (SBDN) in the brain (Albers, 2015; Goodson, 2005; O’Connell and Hofmann, 2011; O’Connell and Hofmann, 2012). In fact, nonapeptide hormones (vasopressin/vasotocin, isotocin/mesotocin, and oxytocin) interact with sex steroids to influence behavior (Goodson and Bass, 2001; Stoop, 2012), making them quintessential behavioral candidate genes.

In sticklebacks, vasotocin peaks during the start of the breeding season in both males and females (Gozdowska et al., 2006). Nesting male sticklebacks have an increase in vasotocin levels in their brains following a mirror (aggression) challenge (Kleszczyńska et al., 2012). The arginine-vasopressin-like (*avpl*) gene showed the greatest overexpression in dominant versus subordinate zebrafish (Filby et al., 2010), further supporting its role in aggressive behavior in teleosts. Vasotocin in adult teleosts is mainly located in the preoptic area (POA) of the hypothalamus (Albers, 2015; Huffman et al., 2012; Kagawa et al., 2016), where it is an active regulator in the hypothalamic-pituitary-adrenal (HPA) axis (Arnett et al., 2016). Therefore, we hypothesized that supplemental expression of arginine vasopressin (*AVP*) within the social decision making network of the stickleback brain would increase aggression.

Exogenous vasotocin administration has been used in many teleosts and other vertebrates to alter behavior, both via intercranial and systemic injection (Goodson and Bass, 2001), and has a dosage based response (Gonçalves and Oliveira, 2011; Moore and Miller, 1983; Santangelo and Bass, 2006). Additionally, given its role as an antidiuretic hormone (Koshimizu et al., 2012), vasotocin produces a rapid physiological response of increased respiration, making its physiological effects quick and non-invasive to monitor. Vasotocin can pass through the blood-brain barrier (Banks et al., 1987; Yaeger et al., 2014). Therefore, unless brain trauma from intercranial injection rendered the technique unsuitable, we expected vasotocin to produce similar effects when administered either by intercranial or intraperitoneal (IP) injection.

Monoamine oxidase, our other candidate gene, has a longstanding association with aggression (Godar et al., 2016), not only in model systems but also in humans (Brunner et al., 1993). Mice with no *MAOA* activity showed increased fearfulness as juveniles and increased aggression in adult males (Cases et al., 1995). Recent work has suggested that *MAOA* allelic variants may be related to the domestication of dogs, a process which included a marked decrease in aggression (Sacco et al., 2017). Teleosts have only one monoamine oxidase gene (*MAO*) as opposed to the two found in mammals (*MAOA* and *MAOB*). Stickleback *MAO* (ENSGACT00000012444.1) and mouse *MAOA* (NP_776101.3) have 68% conservation at the protein level. Despite the low level of conservation, the teleost monoamine oxidase gene has been shown to influence aggression (Freudenberg et al., 2016; Malki et al., 2016; Quadros et al., 2018) and is functionally comparable (Arslan and Edmondson, 2010; Shih et al., 1999). Since a low level of monoamine oxidase is associated with increased aggression, increased expression of *MAOA* was expected to decrease aggression through serotonin turnover.

## Methods

### Animals

Freshwater adult fish were collected in spring to summer in 2016 to 2018 from Putah Creek, CA. Additional F1 fish from crosses generated in 2015 and 2016 and reared in the lab were also used for optimizing the neurosurgical procedure. All fish were housed in the lab in 83 L (107×33×24 cm) group tanks with recirculated freshwater (5 ppm salt). The room was maintained at 18 °C on a 16:8 (L:D) “breeding” photoperiod from April to October and otherwise an 8:16 (L:D) “non-breeding” photoperiod. Fish were one to two years old at the time of their surgery. Males were identified by nuptial coloration (secondary sexual characteristics) and by sexing via PCR (Peichel et al., 2004). Individuals were weighed and measured (standard length from nose to caudal peduncle), and then moved to individual 9.5 L (32×21×19cm) tanks lined with gravel and containing a synthetic plant. Each individual was allowed to acclimate, undisturbed for 3 days prior to any behavioral measurements. All fish only underwent one injection or surgery and were not reused. All animal work was done in compliance with IACUC protocol (#15077 & 18080) at the University of Illinois at Urbana-Champaign.

### Surgical rig

In a ten-minute neurosurgical procedure, a suspension of foreign material (saline, viral construct, or pharmacological agent) was administered to the telencephalon or anterior diencephalon of the brain via transcranial injection. For this procedure, we developed a custom-built surgical rig (Fig. 1). To provide continuous oxygenation and anesthesia to the fish while out of water, a low pressure and flow rate cannula pump was necessary. The complete parts list along with assembly instructions are publicly available through the Open Science Framework (https://osf.io/sgpvm).

**Figure 1.**
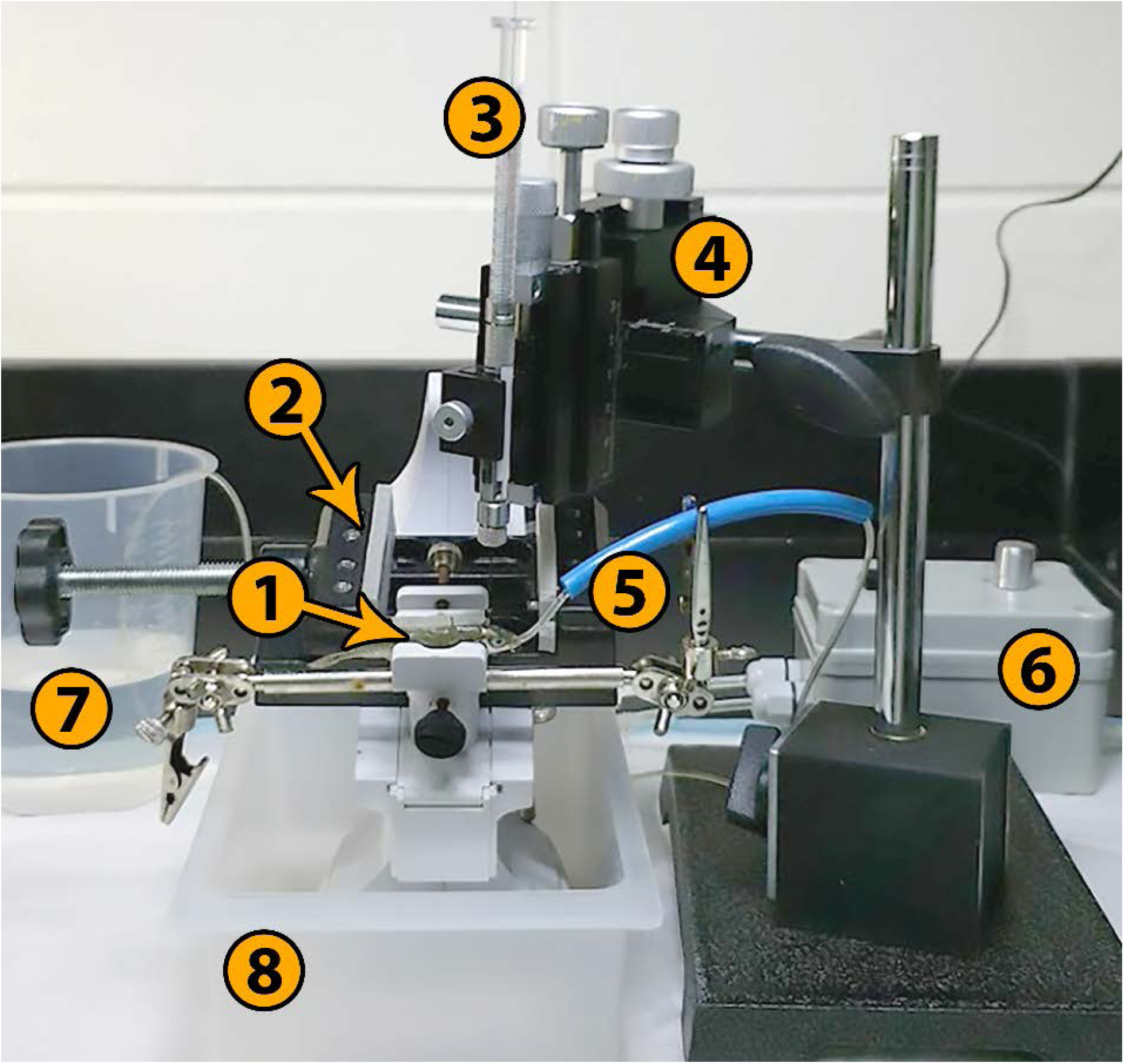
Custom-built surgical rig. 1. Threespine stickleback in padded clamp
2. Alternative padded clamp for larger fish
3. Neuros syringe, 5 μL
4. Three-axis manipulator
5. Oral cannula and guide tube
6. Peristaltic cannula pump, 100 mL/min
7. Pump source reservoir
8. Drip tray

### Anesthesia

Prior to anesthetization, a pre-surgical baseline respiration rate was taken by counting opercular beats per 20 seconds. Initial anesthetization was done in 0.02% buffered MS-222 (Tricane-S, Western Chemical, Fisher) for no more than five minutes (188.4 sec ± 74.0), until movement ceased and the fish was unresponsive. A properly sedated fish had 1) no tail movement, 2) decreased but regular respiration (opercula beating), 3) came to rest on the bottom of the soaking container, and 4) did not respond to touch nor move when removed from the bath.

Anesthetization time was not correlated with any physiological measure (Fig. S1). Fish were rinsed in freshwater (5ppm salt) to remove any residual anesthetic then moved to the surgical rig. In the rig, an oral cannula supplied constant water flow with 0.01% maintenance anesthetic over the gills for the duration of the surgical procedure (233s ± 79, mean ± s.d.). The speed of water delivery was adjusted to each fish to allow a steady low flow rate over the gills.

### Neurosurgical optimizations for direct brain injections

Fish were gently clamped into the surgical rig behind the eyes, keeping the skull firmly in place. The stickleback brain is visualizable through the skull (Fig. 2), allowing injection sites to be selected with moderately high precision. In each injection, the needle, either a 5 μL borosilicate syringe (Hamilton Neuros model 75, #65460-02, Reno, NV) with a 33G (0.210 mm OD) needle or an insulin syringe (BD 328431, Franklin Lakes, NJ) with a 30G (0.337 mm OD), was inserted transcranially through the thinnest portion of the skull. The finer 33G needle had a hard stop set at 2.5mm. Each transcranial injection delivered ~300 nL of liquid at one or three depths to moderate the area transfected, with one depth being the most restrictive. Bilateral transcranial injections delivered a total of ~600 nL. Any notable accumulation of blood was associated with poor outcome.

**Figure 2.**
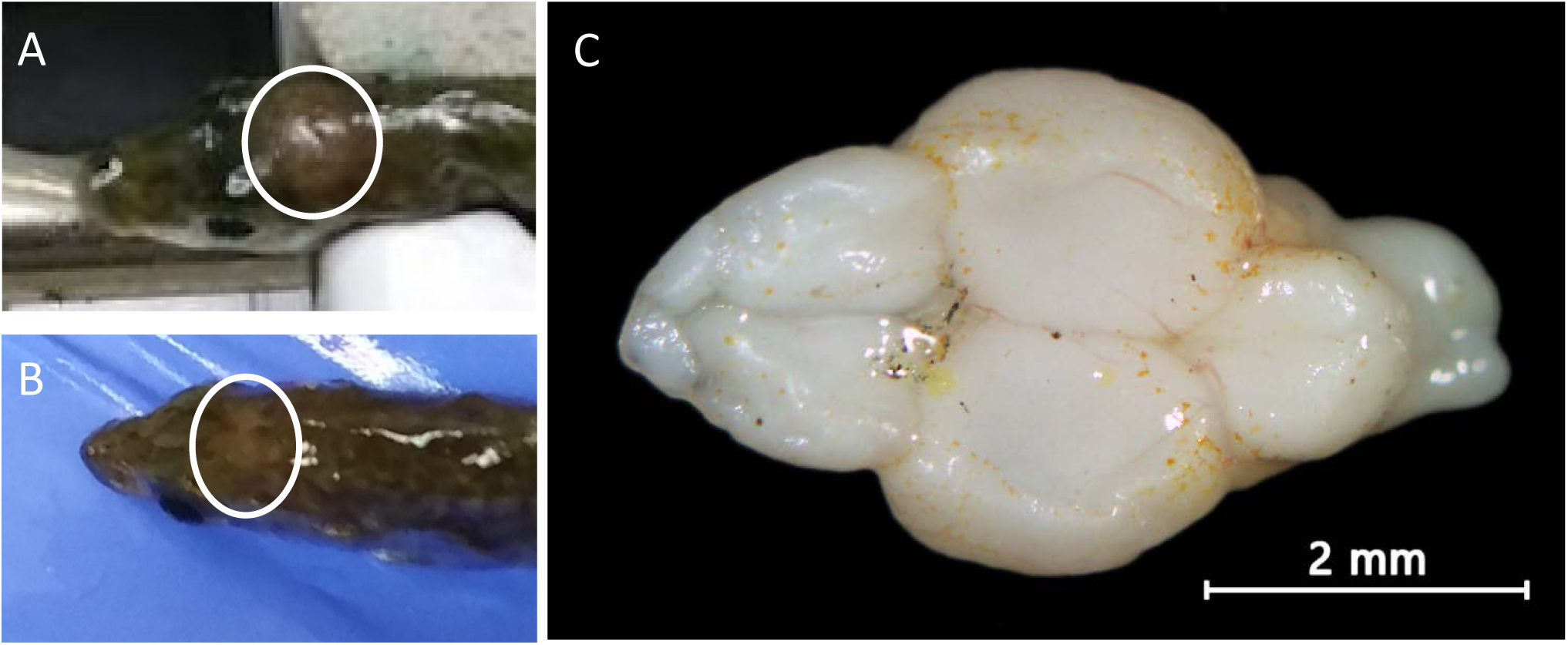
Visualization of the stickleback brain. A & B) Example visualizations of the stickleback brain through the skull. Circled is a lighter area corresponding to the diencephalon. C) The brain of a stickleback with olfactory bulbs (anterior, far left), telencephalon (left), diencephalon and mesencephalon (center), cerebellum (right), and brain stem (posterior, far right). The telencephalon contains much of Social Behavioral Network (amygdala, hippocampus, etc). The preoptic area (POA), hypothalamus, and periaqueductal gray/central gray (PAG) are in the diencephalon.

Following the procedure, fish were returned to their individual tanks and monitored continuously until clear respiration (opercular movement) was seen, typically within 30 seconds. In the case of shallow respiration, forced movement of fresh water over the gills was used to promote survival by manually “swimming” the fish in a submerged figure eight using only forward motion. Respiration rate and the fish’s position in the water column was recorded every 15 minutes for two hours following the injection. Additional checks were performed at three hours and one-day post-injection for all fish. Out of 183 total fish receiving brain injections, 19 did not survive this initial three-hour recovery period; nine did not survive anesthetization and ten were euthanized.

### Neurosurgical procedure changes for viral-mediated transgenesis experiment

In order to accommodate animal biosafety level 2 (ABSL-2) restrictions, fish were transferred into a new tank in the surgery room the morning of the injection. Broad expression throughout both hemispheres of the diencephalon (Fig. 2C) was desired to maximize the probability of producing detectable behavioral changes. Therefore, every fish received two bilateral transcranial injections to the anterior diencephalon of one of the viral constructs delivering a total of ~600 nL of construct across multiple depths from a 5 μL borosilicate syringe with 33G needle. After two days, fish were removed from the ABSL-2 surgical room to individual tanks.

### Pharmacological treatments

Exogenous [Arg8]-Vasotocin (Genscript RP10061, Piscataway, NJ) was administered either directly into the brain via injection using the neurosurgical protocol described above or systemically via intraperitoneal (IP) injection, already a well-established method in sticklebacks, using a 30G (0.312 mm OD) insulin needle. Because behavioral response has been reported to differ in teleosts based on dosage (Gonçalves and Oliveira, 2011; Santangelo and Bass, 2006), a dose-response curve (0.5, 5, and 10 µg per gram body weight) was tested. Manning compound (Bachem H-5350.0001, VWR), a potent V₁ receptor antagonist (anti-vasopressor) was administered systemically via IP injection at a dosage of 3 µg per gram body weight. Both pharmacological agents were freshly diluted on the day of injection from a pre-suspended concentrated stock solution such that all IP injections delivered 10 μL per gram body weight.

Behavioral assays were performed 48 hours prior to pharmacological manipulation for baseline measurements, and then at 30 minutes after IP injection or 2 hours after brain injection. Preliminary saline injections showed that 30 minutes was sufficient for both physiological and behavioral recovery from the IP injection procedure (full data: https://osf.io/v56zt).

### Viral-mediated transgenesis constructs

We elected to use Herpes Simplex 1 (HSV-1) as a viral vector as it was already shown to be effective in transfecting teleosts, where adeno-associated viruses (AAVs) have previously failed (Zou et al., 2014). Additionally, HSV-1 generally has low genotoxicity and no detectable shedding (Bouard et al., 2009) reducing biohazard concerns. Finally, HSV-1 supports larger payloads than AAVs or retroviral vectors such as lentivirus (Howarth et al., 2010; Simonato et al., 2000).

Three promoters in replication deficient HSV-1 were piloted to drive gene expression based on work in zebrafish (Zou et al., 2014) – a long-term promoter (hCMV, *N* = 43, Fig. 3) resulting in fluorescent signal 2-5 weeks after injection, a short-term promoter (mCMV, *N* = 10) with expression between 4 and 7 days post-injection, and a retrograde promoter (hEF1a, *N* = 7) which did not result in a detectable fluorescent signal. Promoters were tested for their ability to drive a fluorescent protein (EGFP, EYFP, GCaMP6f, or mCherry). The long-term promoter (hCMV) was selected as the most useful due to its longer window of effect and was used in the viral-mediated transgenesis experiment.

**Figure 3.**
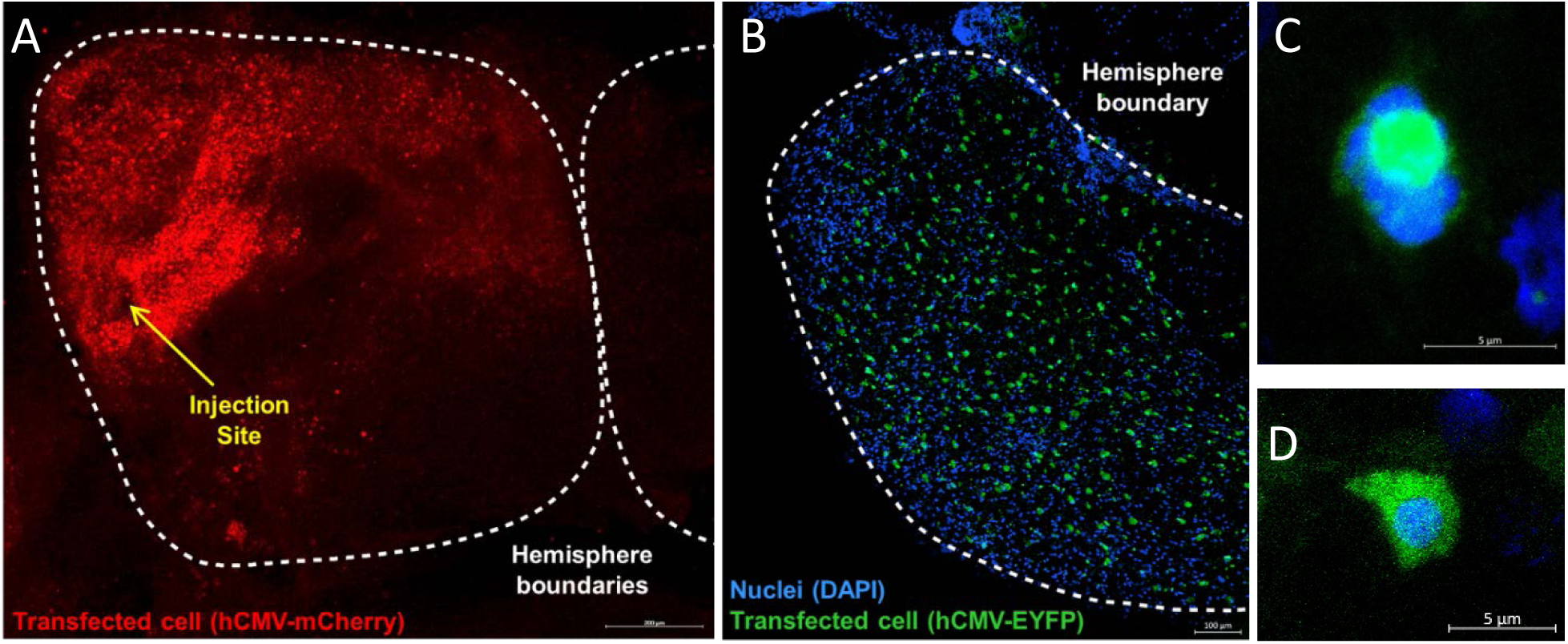
Fluorescent expression resulting from viral-mediated transgenesis. A) Single injection resulting in local expression, limited to a portion of one hemisphere of the telencephalon. B) Broad expression throughout left hemisphere of the diencephalon, typical for injections with delivery at multiple depths. No fluorescence was seen in any saline injected controls. C & D) Successful transfection of entire cells by the long term hCMV-EYFP construct in the lateral left diencephalon three weeks after injection.

Mammalian cDNA ORF clones were used for *AVP* (human, HG17671-UT, NCBI Ref Seq: NM_000490.4, Sino Biological, Beijing, China) and *MAOA* (mouse, MG57436-U, NCBI Ref Seq: NM_173740.3, Sino Biological). These were cloned into the pDONR221 backbone (Epoch Life Science, Missouri City, TX) and then packaged (Gene Delivery Technology Core, Massachusetts General Hospital, Boston, MA) with an IRES-GFP backbone in replication deficient HSV-1. Stock hCMV-EYFP (RN12) was used for control injections. All males were randomly assigned to one of the three constructs. The final viral solutions were used undiluted except for the addition of a trace amount of pigment (brilliant blue FCF or tartrazine, i.e. FD&C Blue No. 1 and Yellow No. 5) to allow the solution to be visualized against the gradations of the syringe. These constructs are episomally expressed; the payload genes, packaged as a plasmid, remain in the cytoplasm and neither integrate into nor replicate with the genome.

### Behavioral assays

All behavioral data were gathered double-blind to treatment (saline, pharmacological agent or transfected gene). Respiration rate was determined prior to the territorial challenge by averaging two separate non-continuous counts of opercular beats per 20 seconds taken within a 5-minute period. This ensured that individual variations due to stress from the researcher’s activity were minimized. Territorial aggression was measured by recording the individual’s response to an intruder confined to a glass flask (derived from Wootton, 1971). The times to orient toward and to first bite at the intruder (TTO and TTB, respectively) were recorded, as well as the total number of bites, charges (lunges), and trips (approaches) during the five minutes following initial orientation (behaviors defined in Wootton, 1971). Intruders (*N* = 9) were 5-10% smaller conspecific males.

During the viral-mediated transgenesis experiment, males’ respiration rate and behavioral response to a territorial challenge were recorded four times (Fig. 4): twice before and twice after injection, respectively considered baseline and transfected. Each focal male except one was confronted by the same intruder during all four territorial challenges. In the exception, the initially paired intruder died between trials two and three and was replaced with a new male of the same length.

**Figure 4.**
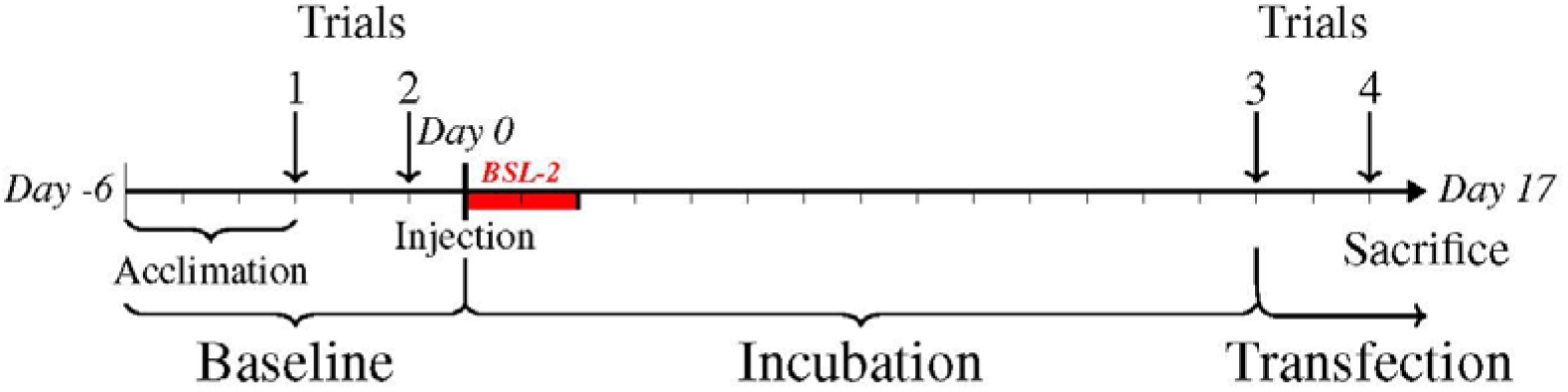
Experimental timeline with the injection of constructs on day 0. Fish were injected with a randomly assigned construct of either an aggression-related gene (*AVP* or *MAOA*) or a control fluorescent protein (*EYFP*). All trials were conducted double-blind to the transfected gene. Each trial had two respiration rate measurements followed by a territorial challenge.

### Histology

Fish were sacrificed via beheading and brains were immediately dissected out. Brains were mounted in glycerol medium on slides with four well iSpacers (SunJin Lab IS018, Hsinchu City, Taiwan). Imaging was performed on the same day as dissection on a Zeiss LSM 710 at the Core Facilities at the Carl R. Woese Institute for Genomic Biology. No fluorescence was seen in any saline injected controls.

### R statistical analysis and data availability

All data are presented as mean ± standard deviation. Boxplots are drawn with a heavy stroke at the median and whiskers of 1.5 IQR beyond the upper and lower quartiles. All data analysis was carried out in RStudio (v1.1.383) with R version 3.5.1. All scripts and data are publicly available on the Open Science Framework (https://osf.io/v56zt) as “Neurosurgical Protocol scripts.R” for the neurosurgical optimization, “AVP pharma treatment behavior.R” and as “Behavioral experiments scripts for release.R” for the viral-mediated transgenesis.

Survival rate differences for the neurosurgical optimization were calculated using the chi-squared function with continuity correction. A nonparametric-compatible repeated-measures ANOVA was done via the `MANOVA.RM` (v0.3.2) package with the ANOVA type statistic (ATS) reported because the assumption of sphericity could not be met for respiration rate over time (Mauchly tests for sphericity = 0.02, *p-value* =1.65e-74). We report the ANOVA-Type Statistic (ATS) and the adjusted degrees of freedom, the latter of which are based on the number of treatment levels, number of observations, and the variance of ranks in each treatment (Shah and Madden, 2004). For interaction effects, we report the recommended 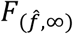 instead of 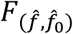 (Noguchi et al., 2012). Post-hoc calculation by time point was done via Wilcoxon rank sum test with continuity correction and the `rcompanion` (v2.2.1) wilcoxonR function. P-values were then adjusted for false discovery rate (fdr method).

Repeatability is reported as ICC3,1 calculated using `Desctool` (v0.99.25) and confirmed with the nonparametric concordance package `nopaco` (v1.0.6). Significance was similar between the ICC and concordance tests. Spearman correlations were calculated using `Hmisc` (v4.1-1). Wilcoxon and Mann-Whitney tests were done with the base `stats` package and effect size was calculated with the `rcompanion` package (v2.2.1). Finally, sample size calculations utilized the `WMWssp` package (0.3.7) with the defaults of 0.05% for two-sided type I error rate and 0.8 power.

## Results

### Neurosurgical optimizations for direct brain injections

Direct transcranial injection with an unbeveled ultrafine needle proved simpler and more effective than other piloted techniques, including craniotomy. Fish generally returned to normal swimming and water column use within 15 minutes after removal of anesthesia. Initial piloting (*N* = 62) revealed respiration rate followed a typical pattern after the operation which we called a recovery curve (Fig. 5). Respiration rate peaked about 30 minutes post-surgery and returned to baseline levels by two hours after the neurosurgical procedure.

**Figure 5.**
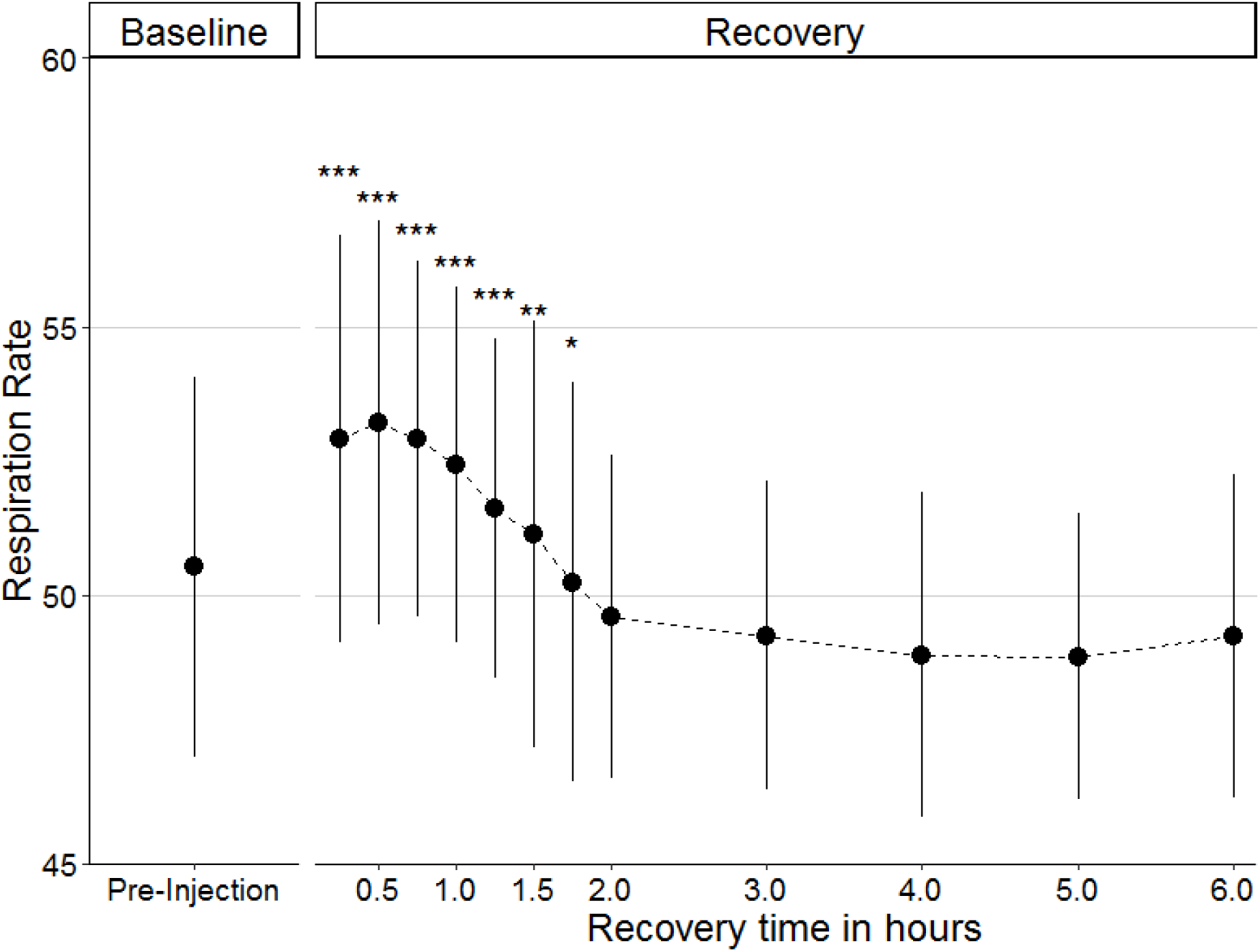
Average recovery curve of respiration rate (opercular beats per 20 seconds) following transcranial brain injections (*N* = 62). Respiration rate returned to baseline levels by two hours post-injection and remained stable following the surgery. * *p* ≤ 0.05; ** *p* ≤ 0.01; *** *p* ≤ 0.001

There was no difference in territorial aggression two hours after brain injection in saline treated controls (*N* = 10), compared to the day before surgery in any aggressive behavior (bites: *Z* = −1.22, *p-value* = 0.21; charges: *Z* = −0.62, *p-value* = 0.54; time to first bite: *Z* = −1.22, *p*-value = 0.22, Fig. 6). Full mating behavior occurred within 3 days for males, determined by nesting behavior and 9 days for females, determined by the presence of eggs.

**Figure 6.**
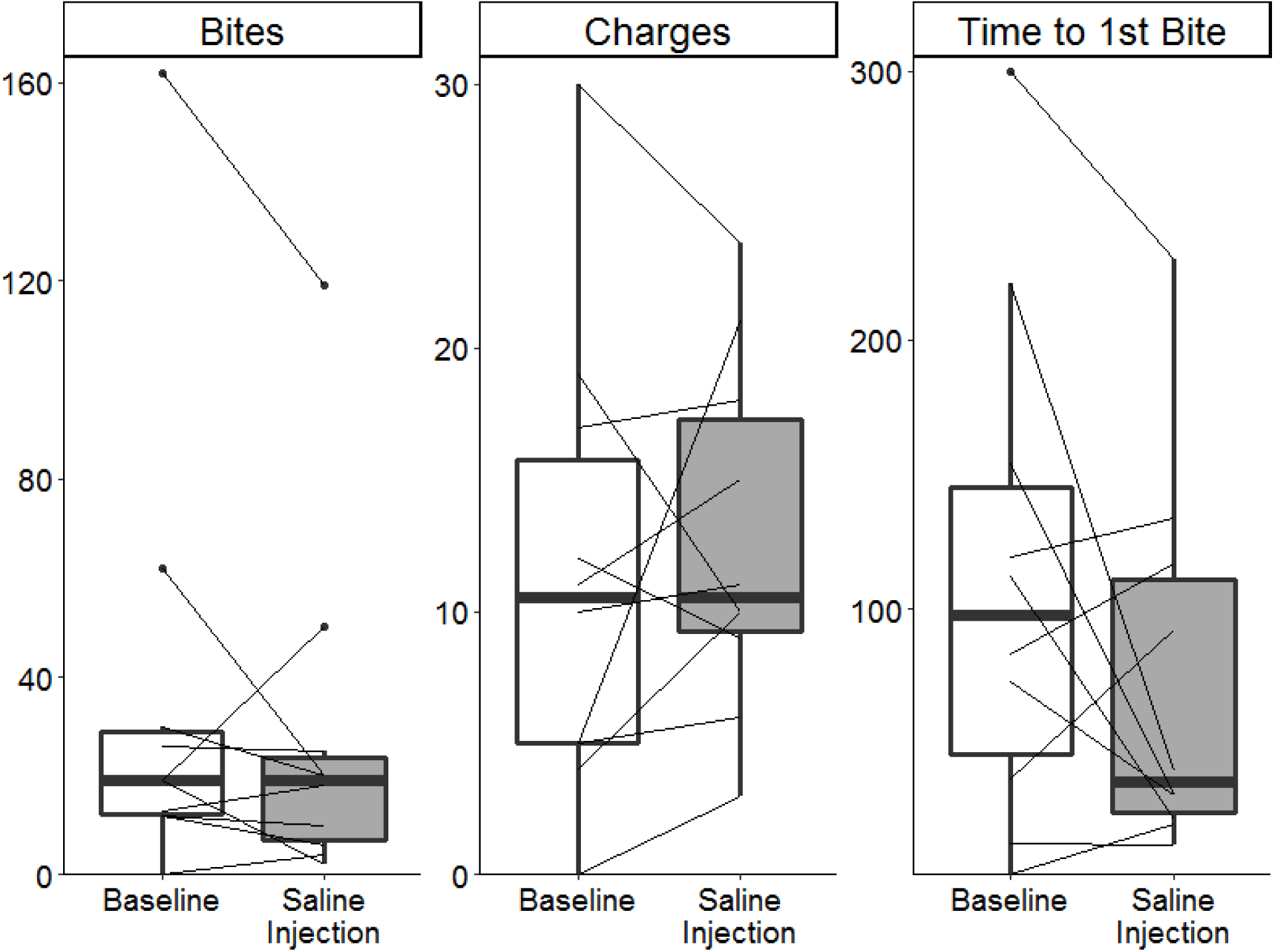
Aggressive behaviors in control fish one day prior to neurosurgery and after direct brain injection of saline (*N* = 10). Each line represents an individual showing their change in behavior following saline injection. No significant change in aggression was seen in any behavior.

Needle diameter influenced survival rate (*χ^2^* (1, *N*_30G_ = 43, *N*_33G_ = 183) = 23.9, *p-value* = 1.02e-6) with a 54% survival rate with the larger 30G needle compared to the 87% survival rate with the finer 33G needle. Additionally, the finer 33G needle resulted in consistently lower respiration rates (*ATS*_1, 2613_ = 6.54, *p-value* = 0.01) during the six-hour window following the neurosurgery than the larger diameter 30G needle. Supplemental oxygenation for up to two days following surgery did not improve survival (*χ^2^* (1, *N*_Extra O2_ = 94, *N*_Normal_ = 89) = 0.02, *p-value* = 0.89) nor recovery (*ATS*_1, 7465_ = 0.94, *p-value* = 0.33, Fig. S3).

### Multiple transcranial injections do not alter survival or recovery

Broad expression required multiple transcranial injections – one into each hemisphere of the brain. Bilateral transcranial injections did not alter survival rates (*χ^2^* (1, *N*_Unilateral_ = 64, *N*_Bilateral_ = 119) = 0.46, *p-value* = 0.50) compared to a unilateral injection. Overall, respiration rate during recovery was higher in fish with bilateral injections (*ATS*_1, 4874_ = 11.61, *p-value* = 0.001) in the three hours following the surgery. However, this comparison was largely confounded by year of capture, with the fish caught in 2018 all receiving two transcranial injections and showing poor health in general. When analyzing only years (2017 & 2019) that received both unilateral and bilateral injections, allowing for direct comparison, there is no difference in respiration rate during recovery between unilateral and bilateral injections (*ATS*_1,312_ = 0.92, *p-value* = 0.34). This suggests that the differences in recovery among years overwhelmed any effect that multiple injections might be having.

### Injection material does not alter survival or recovery

Fish had comparable survival rates regardless of the injected materials (*χ^2^* (3, *N_HSV-1_* = 113, *N_Pharma_* = 22, *N_Saline_* = 48) = 2.30, *p-value* = 0.32). After the surgical technique was refined, the procedure’s survival rate was approximately 90%. Of the 113 fish injected with one of three constructs utilizing replication deficient HSV-1 for transfection, 101 survived. The fish injected with pharmaceutical agents fared similarly, with 20 of 22 fish surviving. Control fish injected with saline fared least well as they were used to initially pilot and refine the surgical technique; they counted 39 survivors among 48 fish, an 81% survival rate.

The time for recovery of fish injected with replication deficient HSV-1 did not differ from that of saline injected controls (*ATS*_3.7, ∞_ = 1.83, *p-value* = 0.12). Additionally, the choice of promoter (hCMV, mCMV, or hEF1a) did not alter survival rate (*χ2* (2) = 1.38, *p-value* = 0.50), nor was there a main effect of promoter (*ATS*_2, 143_ = 0.33, *p-value* = 0.70) on the recovery curve. Finally, the specific gene being expressed had no effect (*χ2* (3) = 2.16, p = 0.54) on survival rates relative to saline injected controls. The recovery rate was also unaffected by the gene expressed (*ATS*_2,267_ = 1.19, *p-value* = 0.31).

### Vasotocin pharmacological treatment

Exogenous vasotocin injection into the brain increased respiration rate rapidly relative to saline injected controls (*ATS*_1,599_ = 8.74, *p-value* = 0.003, Fig. 7). This effect began within 15 minutes and persisted for more than two hours post-injection. The most pronounced difference in respiration rate was between 0.75- to 1.5-hour post-injection. Intraperitoneal injection of exogenous vasotocin also resulted in a rapid increase in respiration rate (Fig. S3). Behaviorally, brain and IP injections of exogenous vasotocin produced parallel results (Table S1), in which only the highest dosage (10 µg per gram body weight) altered the number of charges directed at the intruder.

**Figure 7.**
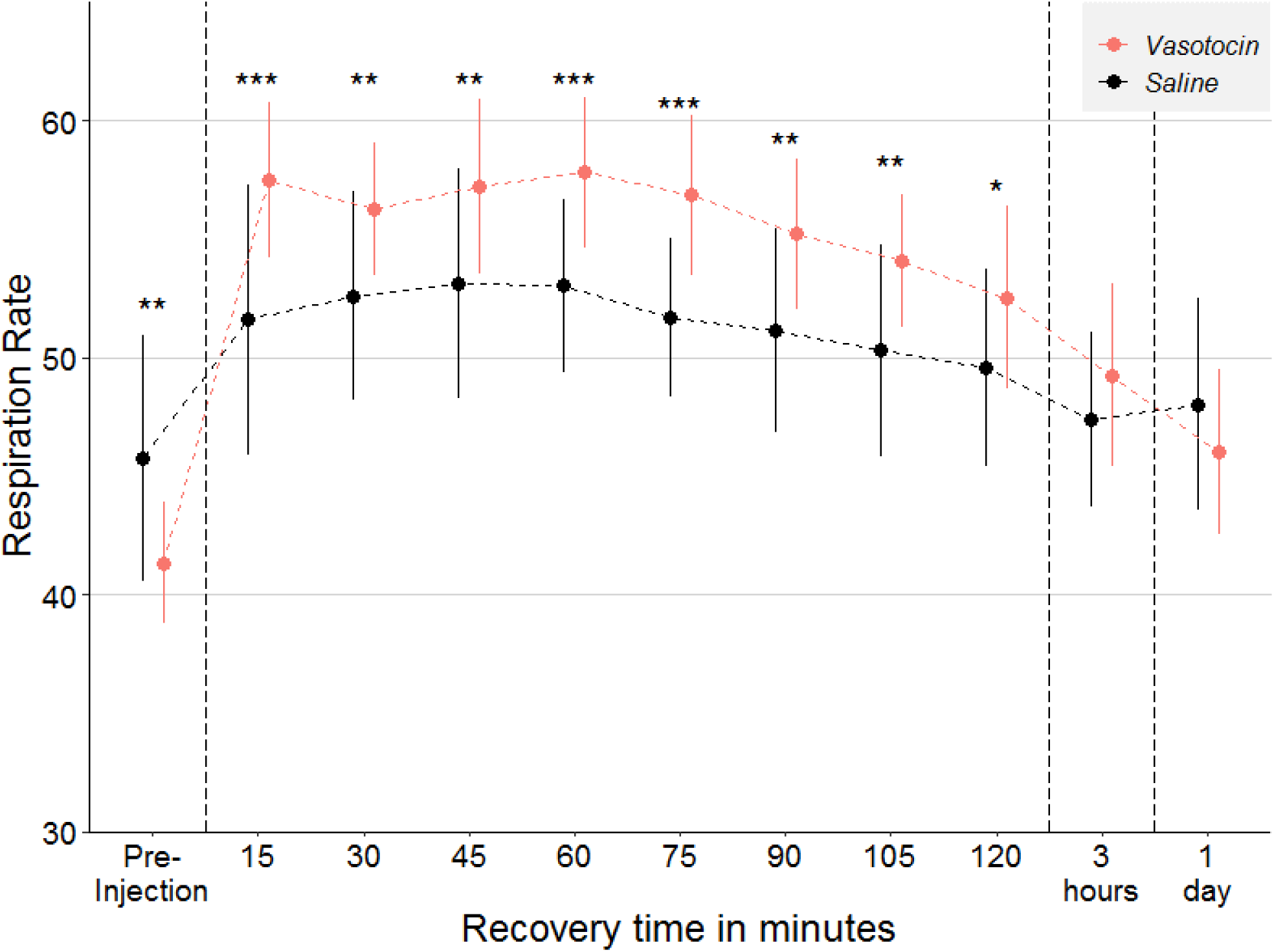
Differences in respiration rate following injection between brain injection of exogenous vasotocin (*N* = 15) and saline injected controls (*N* = 39). Fish injected with vasotocin had an elevated respiration rate compared to saline injected controls for more than two hours following injection. * *p* ≤ 0.05; ** *p* ≤ 0.01; *** *p* ≤ 0.001

### Behavioral repeatability and intercorrelations

Repeatability was analyzed across the two baseline and two transfected trials. Charges, bites, and time to first bite were consistently repeatable, i.e. at both baseline and following transfection but not necessarily between baseline and transfection (Table S2). Aggression measures were generally equally repeatable compared to the physiological measure of respiration rate. Total number of bites and charges were strongly correlated (*r* = 0.69) in the control group (*N* = 16) and following transfection of either *AVP* (*r* = 0.83, *N* = 18) or *MAOA* (*r* = 0.75, *N* = 20). Time to first bite was negatively correlated with total number of bites and charges as well – i.e. fish that bit sooner also attacked more overall. Given its high repeatability and correlation with other behaviors, we focus on charges as a measure of aggressive behavior in subsequent analyses.

### Increased aggression from transfection of MAOA or AVP but not in controls

Aggressive behavior (charges) increased in fish transfected with either *AVP* (*N* = 18) or *MAOA* (*N* = 20). Injection of the control construct (*EYFP*, *N* = 16) did not significantly alter the number of charges (paired Wilcoxon signed rank test: *Z* = −0.01, *p-value* = 0.50) relative to baseline. Due to the high inter-individual variation (σ^2^*_EYFP_* = 103, σ^2^*_AVP_* = 113, σ^2^*_MAOA_* = 100), there was no significant difference between either *AVP-* or *MAOA-*injected fish compared to controls in charges following transfection (Mann-Whitney test: *AVP*-*EYFP*: *Z* = −0.41, *p-value* = 0.34, *MAOA*-*EYFP*: *Z* = −0.7, *p-value* = 0.48). No increase in aggression was seen in the saline injected controls neurosurgical optimization experiments (Fig. 6). Thus, only injection with either of two aggression-related candidate genes resulted in altered behavior and only specifically in charges (Fig. S4).

*AVP* had a large effect on the number of charges (paired Wilcoxon signed rank test: *rs* = 0.79, *Z* = −3.07, *p-value* = 0.001) with 16 of the 18 individuals increasing the average number of charges compared to their baseline. In magnitude, this represented an almost 100% increase in average number of charges, from 9.7 (s.d. = 5.1) at baseline to 18.8 (s.d. = 10.6) following transfection.

Transfection with *MAOA* also caused a large increase in the average number of charges (paired Wilcoxon signed rank test: *rs* = 0.53, *Z* = −2.10, *p-value* = 0.018) relative to baseline. However, the effect of *MAOA* was less drastic than that of *AVP* and had more variation in individual response (Fig. 8), with 13 of 20 individuals increasing their average number of charges. Despite this, *MAOA* still resulted in an approximately 50% increase from 12.1 (s.d. = 9.7) charges at baseline to 19.1 (s.d. = 10.0) following transfection.

**Figure 8.**
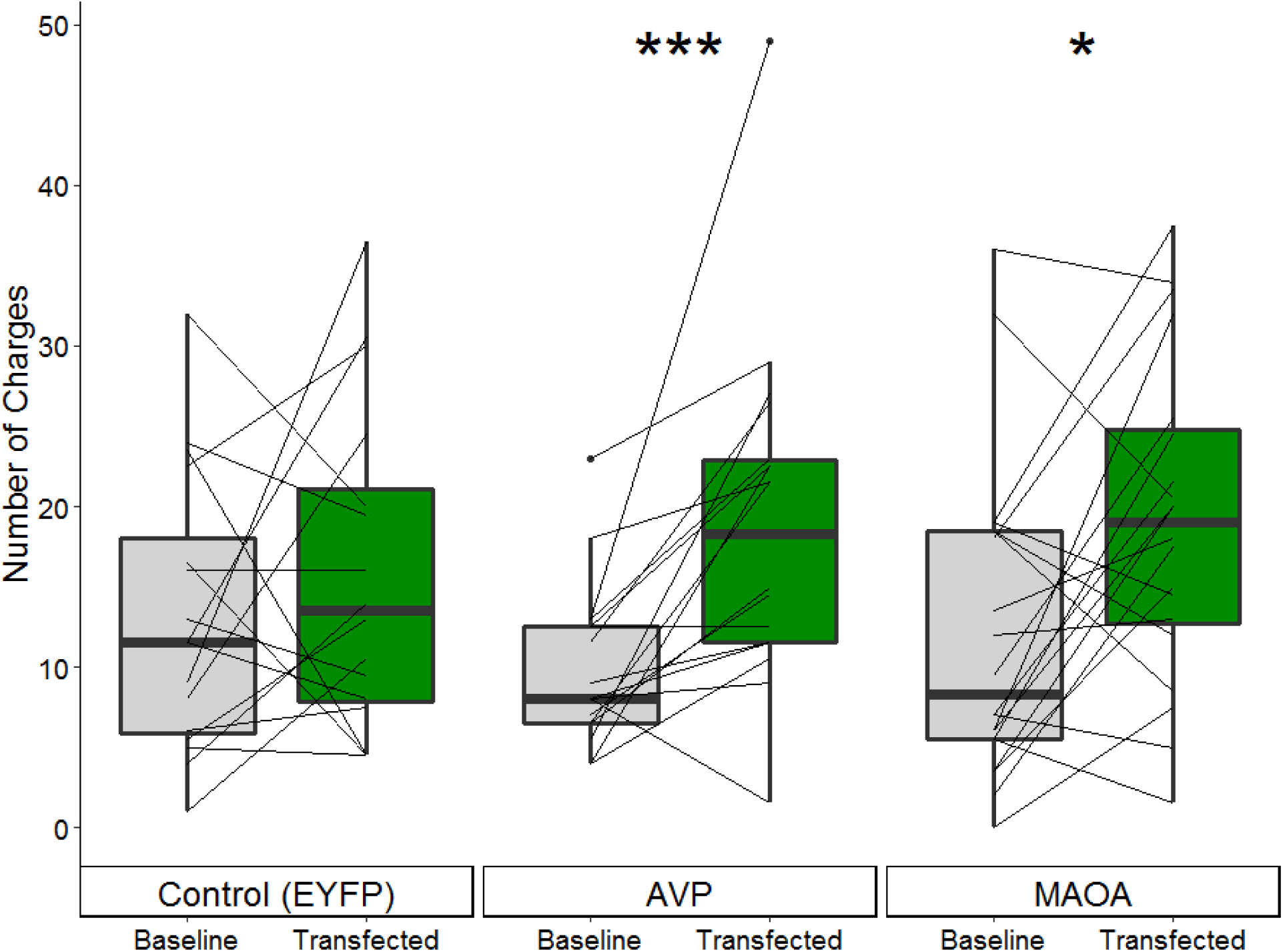
Number of charges (averaged across the two trials) before and after transfection for each construct. Each line represents an individual showing their change in behavior following transfection of the gene of interest. Transfection with *AVP* resulted in a substantial and consistent increase in the number of charges; note that only one individual exhibited decreased charging behavior. Transfection with *MAOA* resulted in an increase of large effect size in charges, although there was more variation in individual response. * *p* ≤ 0.05; ** *p* ≤ 0.01; *** *p* ≤ 0.001

### MAOA decreased respiration rate

Only the *MAOA* construct altered respiration rate (Fig. 9). Compared to baseline, *MAOA* strongly and significantly (*N* = 20, *rs* = 0.85, *Z* = −3.62, *p-value* = 0.0001) lowered respiration rate, with 19 of the 20 individuals experiencing a decrease in resting respiration rate. They dropped from an average of 40.8 (s.d. = 1.9) to 38.1 (s.d. = 2.0) breaths per 20 seconds.

**Figure 9.**
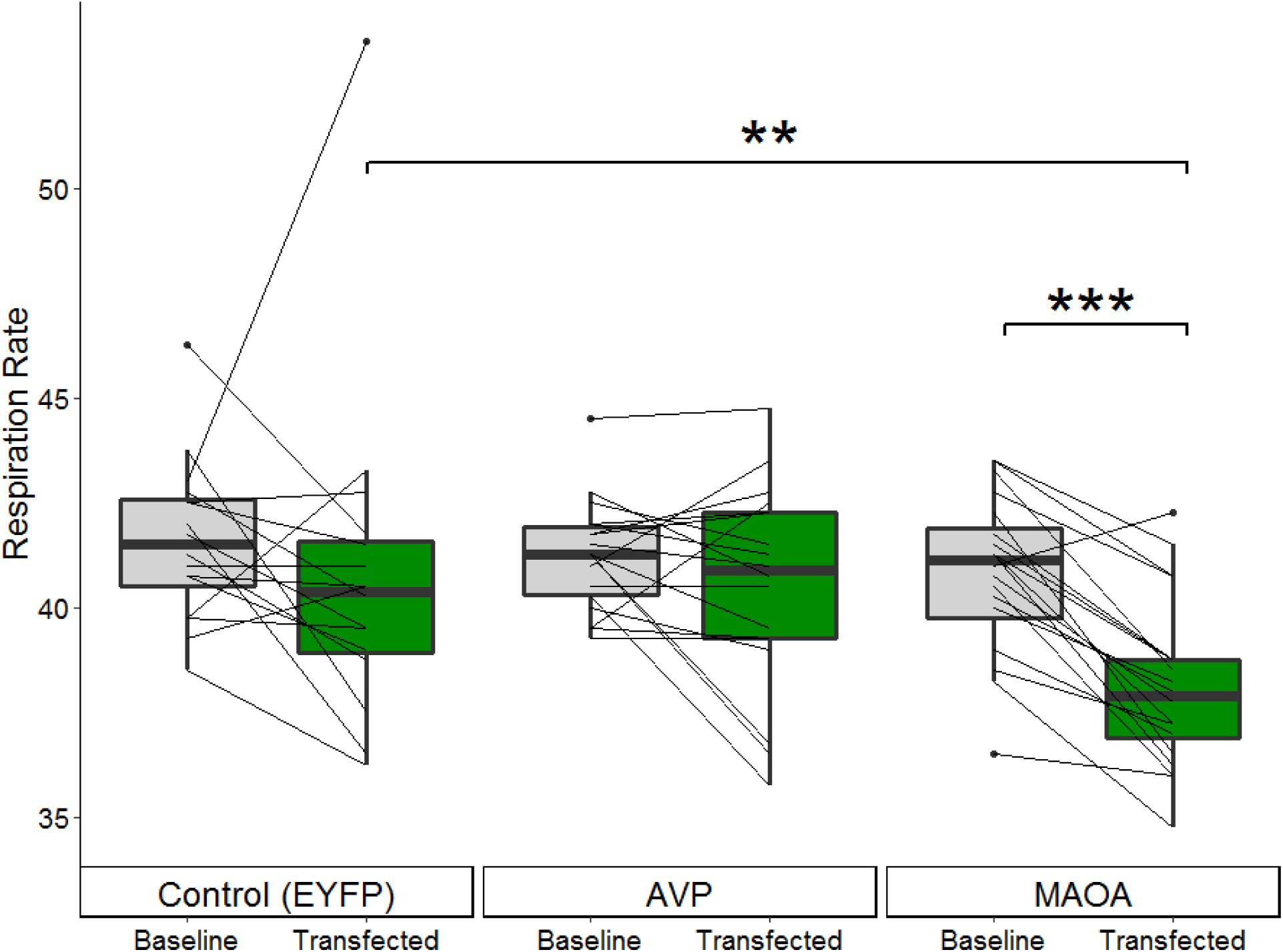
Respiration rate (opercular beats per 20 seconds) for the three constructs. Each line represents an individual showing their respiration rate (averaged across the two trials) before and after transfection. Only *MAOA* altered respiration rate; a drastic decrease compared to both baseline (within-subject comparison, *N* = 20) and to the control group (*N* = 16). * *p* ≤ 0.05; ** *p* ≤ 0.01; *** *p* ≤ 0.001

Additionally, when comparing the respiration rates between the fish transfected with *MAOA* and the *EYFP* controls (*N* = 16, *mean* = 40.8, s.d. = 3.9), the decrease was still significant, though reduced to a moderate effect size (Mann-Whitney test: *rs* = 0.44, *Z* = −2.65, *p-value* = 0.009). There was not significant change in respiration rate compared to baseline due to either the control *EYFP* (*Z* = −1.11, *p-value* = 0.13) or *AVP* (*Z* = −0.92, *p-value* = 0.18) constructs.

## Discussion

### Neurosurgical optimizations for minimally invasive direct brain injections

We present a new method for direct injection of transgenic or pharmaceutical material into the brains of the small teleost fish threespined stickleback. Developing a minimally invasive neurosurgical protocol required 1) refining the anesthesia process, 2) building a custom surgical rig, and 3) determining the normal recovery pattern allowing us to clearly identify warning signs of failure to thrive. Our surgical rig and optimized anesthetization methods (Neiffer and Stamper, 2009; Sladky and Clarke, 2016) resulted in high (90%) survival rates and quick behavioral recovery. Mating behavior also recovered promptly: males completed nests at three days post-surgery, and females were gravid at nine days – suggesting almost no delay in the egg development time (Baker et al., 2008) after losing any ripe eggs to clamping during surgery.

Establishing a typical recovery curve (Fig. 5) allowed us to identify post-surgical warning signs of failure to thrive. Behavioral manifestations of discomfort or problems included listing (>45º off central axis), assuming a nose-up position, and loss of positional control (twirling). The presence of any of these markers for greater than an hour suggested a poor prognosis and thus we recommend euthanasia. Survival to 24 hours indicated a successful procedure, as 23 out of 24 fish injected with the larger 30G needle and 159 of 160 fish injected with the smaller 33G needle survived to one week. Thus, this minimally invasive neurosurgical method is quite reliable.

### Pharmacological Manipulation: bypassing the blood-brain barrier

Exogenous vasotocin administered directly to the brain produced physiological and behavioral responses mirrored in fish receiving vasotocin through IP injections (Figs. 7, S3, Table S1). These pharmacological results were similar to those seen in other fish (Filby et al., 2010; Lema and Nevitt, 2004; Santangelo and Bass, 2006). This indicates that brain injection is now a feasible delivery route for drugs that do not pass through the blood brain barrier (Cook et al., 2009) in sticklebacks. Ultimately, similar changes in respiration and aggression suggest that the recovery period from the brain injection does not mask even rapid onset pharmaceutical effects, like those due to vasopressin (Stark et al., 1989).

In this experiment, similar physiological effects were produced throughout the dose-response curve (Fig. S3). In conjunction, although behavioral effects were produced only at the highest dosage, they were similar for both routes of administration (Table S1). These consistencies suggest that the pharmacological manipulation did in fact work.

### Viral Mediated Transgenesis: Genetic underpinnings of territorial aggression

To demonstrate the practicality of viral-mediated transgenesis to examine candidate genes’ contribution to behavior, we looked for behavioral changes from two aggression related candidate genes. It is well established that both vasopressin (*AVP*) and monoamine oxidase (*MAOA*) influence aggression (Goodson, 2013). Using a ubiquitous promoter resulted in altered territorial aggression specifically for fish receiving either one of the candidate genes related to aggression (Fig. 8), but not for a fluorescent protein (Fig. 8) or following saline brain injection (Fig. 6). The effect following transfection of *AVP* was consistent, with 16 of the 18 fish experiencing an increase in aggression, one remaining constant and only one decreasing aggression.

The effect of *AVP* transfection was of stronger magnitude (*rs =* 0.79) than the effect of pharmacological manipulation of vasotocin (*r_Brain_ =* 0.66, *rIP =* 0.67). This is likely due to the broad expression throughout the diencephalon. It is highly likely that our transgenic procedure resulted in ectopic expression as vasotocin is typically only produced in the preoptic area (POA) and ventral hypothalamus (VH) (Kagawa et al., 2016; Maruska et al., 2019). Maximal expression was desired to increase the likelihood of transfection and of producing behavior changes through supraphysiological levels of vasotocin signaling. It will be fruitful for future studies to examine the consequences of altered expression in key brain areas and cell types, as discussed below.

Multiple pieces of evidence indicate that we achieved successful transgenesis, in addition to the consistent changes in both behavioral (Fig. 8) and physiological (Fig. 9) phenotypes following transfection. The most visible, literally, is the presence of novel fluorescent proteins (Fig. 3). Additionally, transcripts specifically of the injected mammalian *MAOA* homolog remained detectable through qPCR for up to 4 weeks after transfection (results not shown).

Our finding that transfection of *MAOA* increased aggression is consistent with a decrease in serotonin, which is enzymatically cleaved by monoamine oxidase. Indeed, the clear and unambiguous decrease in respiration rate we observed (Fig. 9) is strong evidence of *MAOA* functioning as expected physiologically. Respiration rate correlates positively with serotonin and norepinephrine concentration (Hodges and Richerson, 2008; Whelan and Young, 1953); monoamine oxidase enzymatically lowers levels of both neurotransmitters. Additionally, trout monoamine oxidase has been found to be equivalently effective to human monoamine oxidase in metabolizing 5-HT and PEA (Shih et al., 1999), making it unlikely that the increase in aggression is an off-target effect of using mammalian *MAOA*. Further characterization of anxiety levels following transfection, pharmacological rescue (Godar et al., 2014), and quantification of the downstream neurotransmitters remain as potential avenues to a better mechanistic understanding of this result.

We found that construct selection is not limited to native genes; widely available mammalian plasmids successfully altered behavior. Therefore, this method is therefore accessible to a broad array of users (Dickinson et al., 2020), a feature especially important in a system with roots in ethology. Furthermore, this application of viral-mediated transfection enables within-subject experimental designs, reducing sample sizes required for the same statistical power and making behavioral experiments viable, as detailed below. Finally, rapid behavioral recovery makes viral-mediated transgenesis a viable technique for direct manipulation of candidate genes.

### Future applications of viral-mediated transgenesis in stickleback

While sticklebacks are a non-traditional genetic model system, they are one of the best studied behavioral systems, with well described intra-specific variation in aggression, antipredator behavior, and parental care (Fang et al., 2018; Hendry et al., 2013; Huntingford and Ruiz-Gomez, 2009). Previous studies have identified hundreds of genes that are differentially expressed in the brain in response to a social interaction (Bell et al., 2016; Bukhari et al., 2017; Greenwood and Peichel, 2015; Greenwood et al., 2013; Laine et al., 2012; Mommer and Bell, 2014; Sanogo et al., 2011). However, most of these studies are correlative, and thus the direction of the causal relationship – much less the mechanisms by which changes in gene expression underlie behavior – are still not clear. This method will allow future studies to rigorously test how these genes contribute to behaviors, from detailed mechanistic analyses at the protein level to studies of individual behavior at the organismal level.

A more detailed, cross-population study of vasotocin in stickleback would present an ideal opportunity to investigate the evolutionary constraints or trade-offs between behavior and physiology for pleiotropic genes. In addition to being associated with behavior, vasotocin plays a key role in osmoregulation via the AVP V2 receptors. Vasotocin anatomy and aggression differed in concordantly with salinity and osmoregulation challenges in pupfish (Lema, 2006). A more nuanced examination is possible in stickleback as there are numerous freshwater and several anadromous populations that are independently evolved from ancestral-like marine population. In addition, population differences in aggression (Bell, 2005; Dingemanse et al., 2007; Keagy et al., 2016) and osmoregulation (Hasan et al., 2017) have already been documented in sticklebacks. However, an examination of the integration of these two evolutionary concerns has not yet been undertaken, despite a relationship between aggression and kidney size having already been discovered in stickleback bred for extremes of territorial aggression (Bakker, 1986). This makes stickleback uniquely suited system to address the relationship of physiological ecology, anatomy, and social behavior (Goodson, 2013). Other possible physiological signaling pathways amendable to genome editing in sticklebacks have also been proposed (Dalziel et al., 2012; Hohenlohe et al., 2010; Ishikawa and Kitano, 2020; Jones et al., 2012; Shimada et al., 2011).

### Behavioral response to vasopressin/vasotocin

In every case of vasotocin signaling manipulation that we tested – pharmacological IP inhibition (Manning compound) or supplementation, exogenous brain injection, and transfection – number of charges at the intruder was the main responding aggressive behavior (Fig. 8, Table S2). Charges as the main responding behavior matches the effects of vasotocin seen in pupfishes (Lema and Nevitt, 2004). This is of interest as charging marks voluntary initiation of aggression, while biting is an escalation based off intruder response (van Iersel, 1953). However, depending on the type of manipulation we saw opposing effects on the number of charges – pharmacological supplementation of exogenous vasotocin resulted in a decrease while transfection with *AVP* resulted an increase. As previously discussed, there is reason to hold high confidence in both pharmacological treatments and the transfection method, despite the varying directions of effect.

These techniques have drastically different timings and durations of the increase in vasotocin, potentially explaining the differing directions of behavioral changes. The pharmacological manipulation involved a single dose and a behavioral assay after two hours, while the transgenesis experiment involved increasing exposure throughout a two-week transfection incubation period, with two behavioral assays over 3 days (Fig. 4). Vasopressin has an extremely short half-life of less than a minute (Stark et al., 1989), making pharmacological manipulation rapid but ephemeral. In contrast, transfection is much longer lasting, allowing for long-term stabilization of the HPA axis and commensurately altered behavioral response.

Additionally, vasotocin signaling is mainly constrained by receptor type and location (Albers, 2015; Goodson, 2013; Huffman et al., 2012) further emphasizing the potential of long-term homeostasis to influence the behavioral outcome. To mimic this with pharmacological manipulation, future work should use an implanted cannula in the brain to deliver repeated low doses of exogenous vasotocin. This would allow a more direct comparison between treatment methods clarifying if the opposing effects of pharmacological manipulation and transgenesis are due to homeostatic balancing from long-term exposure or a potential side effect from an immune response to the injection of small molecules.

### Viral-mediated transgenesis allows statistically powerful repeated measures design

Complex phenotypes that emerge at the whole organism level, such as behaviors like aggression, are difficult to assay due to their subtleties and time-intensive screening. Social behaviors are influenced by many genes of small effect (Spencer et al., 2009; Wahlsten, 2012) and social psychology generally has smaller effect sizes (*r*) relative to other psychological sub-disciplines (Schäfer and Schwarz, 2019). Indeed, the neuroscience, psychiatry, psychology, and behavioral ecology fields are plagued with reports of overestimates of effect sizes (Button et al., 2013; Fanelli and Ioannidis, 2013; Forstmeier and Schielzeth, 2011; Schäfer and Schwarz, 2019). Additionally, behavior in natural populations tends to have high inter-individual variation, further reducing statistical power (Taborsky, 2010). For sticklebacks, who have a generation time of approximately one year, the traditional approach of breeding a stable transgenic line is not always practical. Here, by using within-subject design, we successfully examined two behaviorally relevant genes for effects on aggression in wild-caught fish.

By using repeated measures on the same fish before and after transfection, we were able to drastically reduce the necessary sample size needed to detect significant changes in behavior (Table 1). In this study we found large effect sizes for both behavior and respiration rate, a typical physiological measure. However, variation following transfection with *MAOA* was about 25 times larger for charging behavior (*σ*^2^ *=* 99.9) compared to respiration rate (*σ*^2^ *=* 3.9). A between group comparison would have required an impractical sample size of as many as 300 fish to detect the difference in charges, even though these genes have a large magnitude (*rs* > 0.5) of effect on behavior. However, by using these methods we were able to reduce the sample size down to merely 20 fish, a far more manageable number. Thus, viral-mediated transgenesis enables the study of genetic effects on natural behavior in wild-caught animals, because it makes possible a repeated measures design comparing within the same individuals, increasing sensitivity.

**Table 1.** Benefit of within-subject experimental design on sample size for behavioral genetic studies. Necessary sample sizes assume a statistical power of 0.8 and are based on the effect sizes observed in this study – i.e. 8 out of 10 experiments using samples this large will detect the difference. Viral-mediated transgenesis makes possible a repeated measures design comparing within the same individuals, increasing sensitivity. Note that *AVP* and *MAOA* are known to have a large effect on behavior, so this should be viewed a minimum for future candidate genes being screened for behavioral phenotypes.

|  | **Behavior**<br>**(Charges)** |  | **Physiology**<br>**(Respiration)** |
| --- | --- | --- | --- |
|  | *AVP* | *MAOA* | *MAOA* |
| Necessary sample size<br>(between groups, vs. control) | 277 | 187 | 35 |
| Actual sample size | 18 | 20 | 20 |
| Effect size ($rs$) | 0.79 | 0.53 | 0.85 |
| Variance ($\sigma^2$, post-expression) | 113 | 99.9 | 3.9 |

### Conclusion: examining genetic contributions to behavior

Viral-mediated transgenesis is a method to alter a gene’s expression in a specific location or during a controlled timeframe. Refinements to both the stereotactic procedure and the design of constructs promise improvements in the specificity of targeted brain areas and cell types, allowing manipulations of gene expression with great precision. We successfully used multiple promoters to drive expression, tailoring expression profiles through time. While we used ubiquitous promoters with differing timings, cell-specific targeting can be done by using alternate promoters (see Ingusci et al., 2019). There is also tremendous potential for this method to be used in combination with cell type specific promoters that, for example, target astrocytes (*GFAP*), glutamatergic (*vGLUT, PAG*), GABAergic (*GAD*), dopaminergic (*TH*) or prolactin (*PRL*) neurons.

When combined with pharmacological manipulation, DREDDs (designer receptors exclusively activated by designer drugs, reviewed in Roth, 2016) can target neural signaling with extremely precise timing. This method is already being used to identify neural circuits via chemical silencing or activation of receptors including those that are serotoninergic (Ingusci et al., 2019). Finally, viral-mediated transgenesis also lays the groundwork for optogenetics, although there are still engineering challenges to design light, tether-free setups that do not interfere with complex behaviors in fish that weigh less than 2 grams.

Here we present a minimally invasive neurosurgical procedure for sticklebacks that enables viral-mediated transgenesis in the brain as well as pharmaceutical delivery directly to the brain. This method for viral-mediated transgenesis allows for a more direct examination of the genetic mechanisms underlying behavior in wild-caught animals from natural populations. It is flexible, fast, and allows us to compare individual behavior before and after transgenesis, maximizing statistical power. It further enhances the growing molecular toolkit in threespine stickleback, a classic ethological system. Overall, our experimental results show that viral-mediated transgenesis is a promising method for testing the function of candidate genes in this system. This approach has already proved essential in the functional testing of genes related to behavior (Gallant and O’Connell, 2020; London, 2020; Simonato et al., 2000) and in the dissection of neural circuits (Luo et al., 2008) in other organisms.

## Acknowledgements

Thanks to the Bell lab, especially Colby Behrens, Miles Bensky, Severin Odland, Christian Zielinski, Brianna Bowman, and Rachael Kirchschlager who assisted with neurosurgeries. Colby Behrens also provided the stickleback brain image (Fig. 2C). Neurosurgery consultations provided by Dr. Helen Valentine, DVM, MS, DACLAM & Dr. Jennifer Criley, DVM, DACLAM. Viral packaging was provided by Dr. Rachael Neve of the Massachusetts General Hospital’s Gene Delivery Technology Core. Dr. Rhanor Gillette and Dr. Gene Robinson provided equipment. Brian James assisted with surgical apparatus design and manuscript editing. This work was supported by NSF Edge Grant # 1645170 awarded to Dr. Daniel Bolnick, and co-PIs Alison Bell, Michael White, Craig Miller, and Kathryn Milligan-Myhre.

## Supplemental Figures and Tables

**Supplemental Figure 1.**
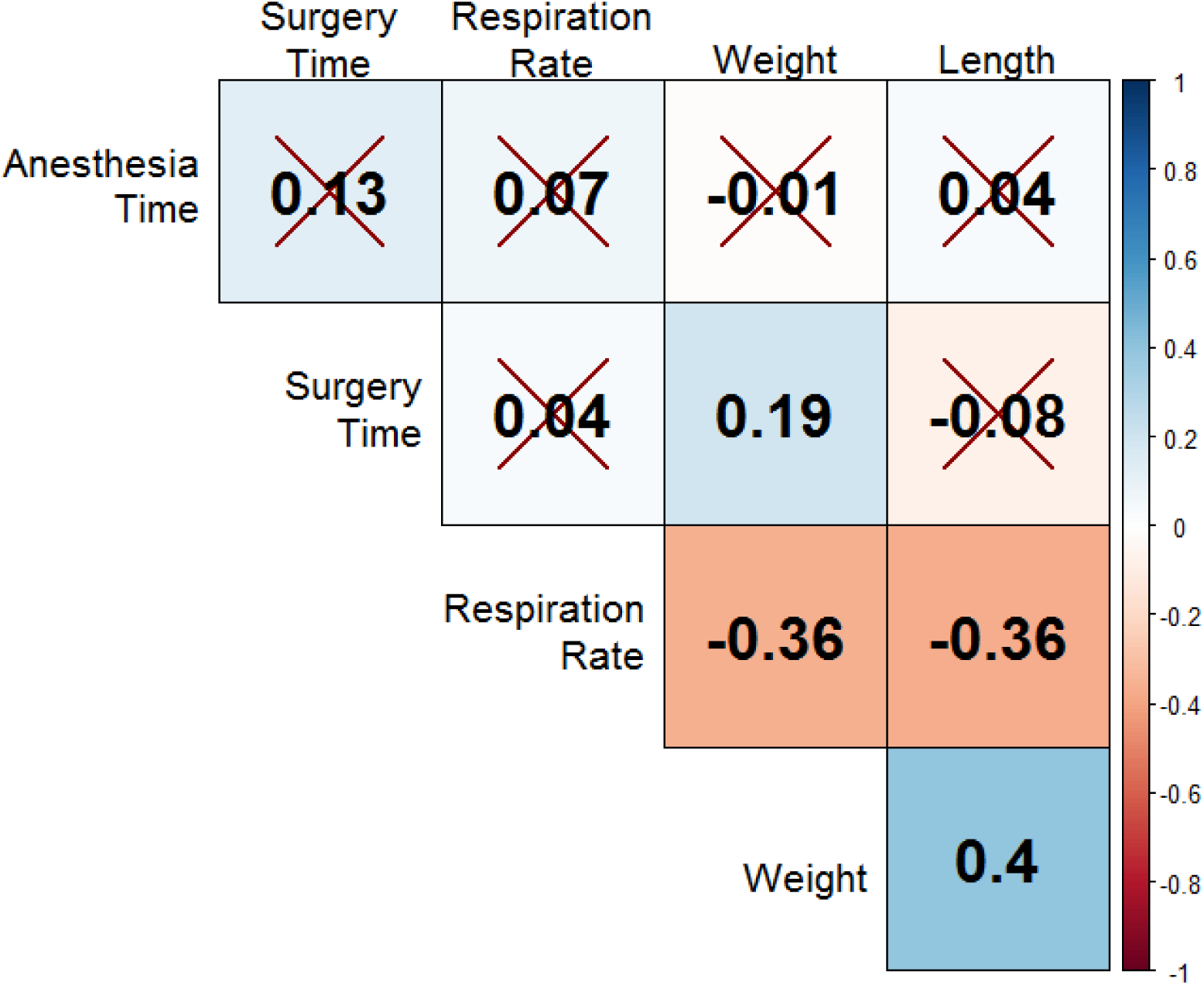
Spearman correlations between time to anesthetization (*N* = 148), time for neurosurgical procedure (*N* = 124), respiration rate prior to surgery (*N* = 153), weight (*N* = 157) and length (*N* = 87). Anesthetization time was not correlated with any other measure. Larger fish had slower respiration rates and it took longer to perform the surgery on heavier fish, in large part due to increased care in clamping. Color indicates direction and strength of correlation. Non-significant correlations have a cross-out.

**Supplemental Figure 2.**
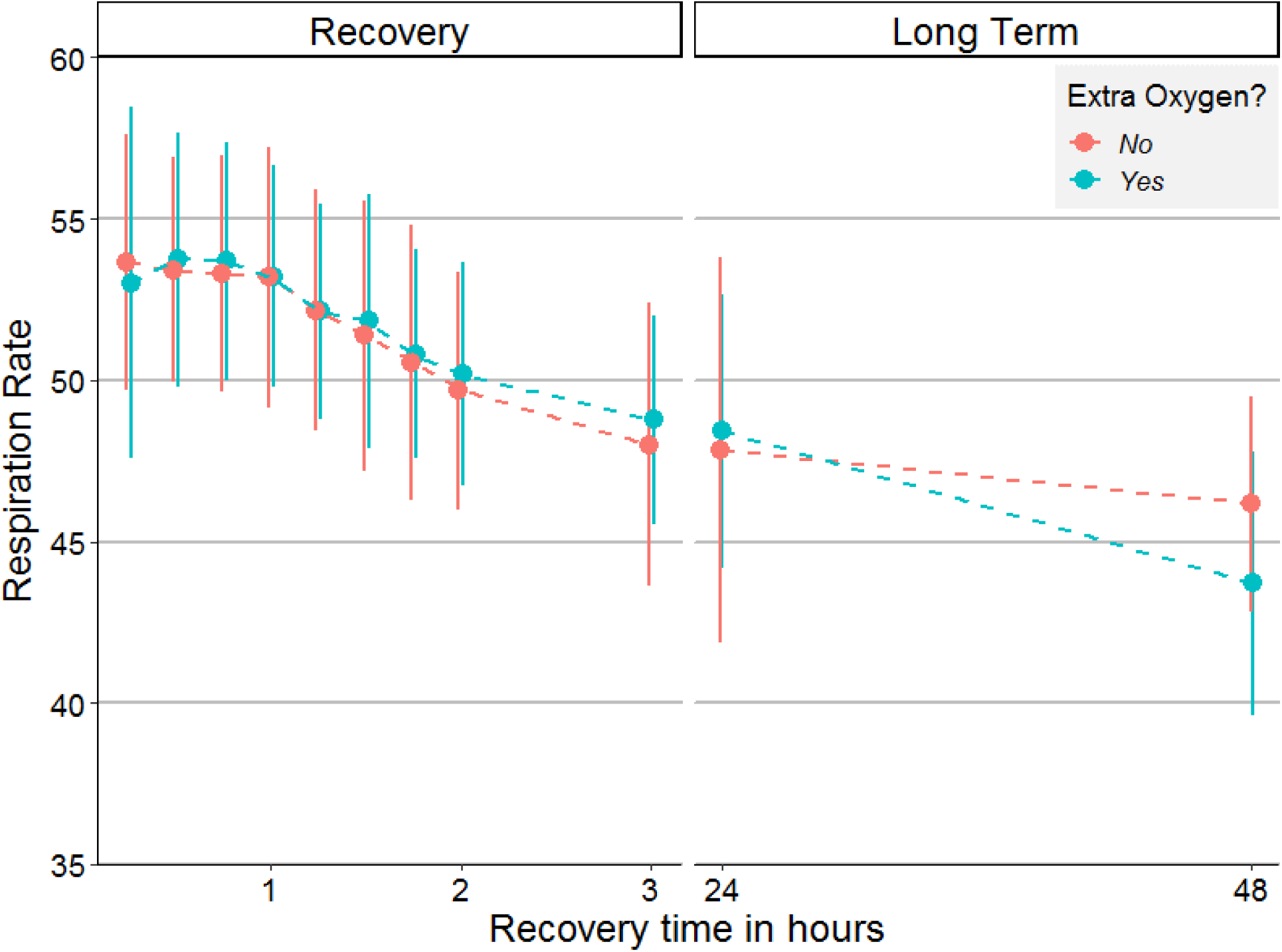
Respiration rate (opercular beats per 20s) following injection either with (*N* = 83) or without (*N* = 77) supplemental oxygenation for one day following the neurosurgical procedure. Supplemental oxygenation did not significantly improve recovery rates.

**Supplemental Figure 3.**
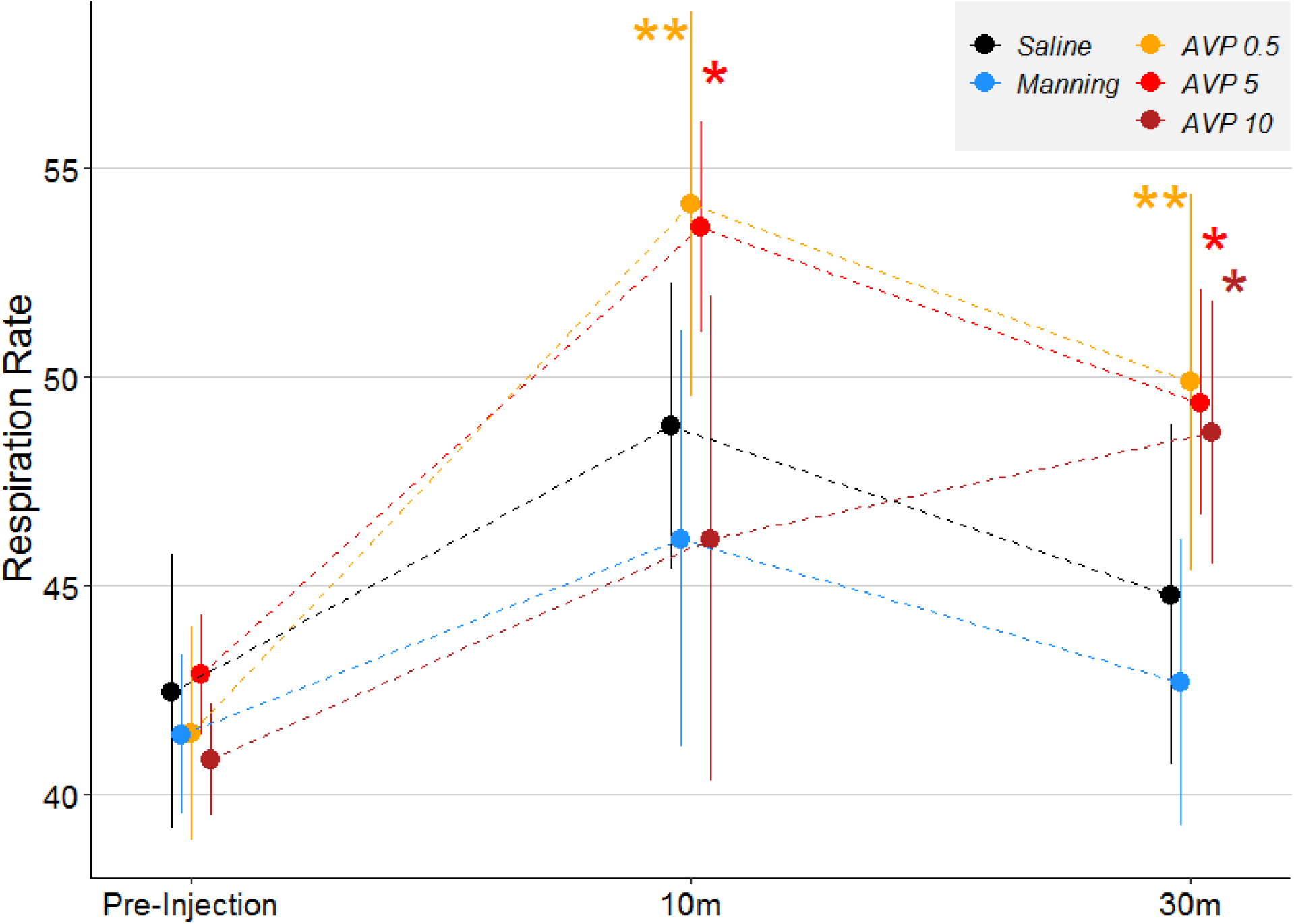
Differences in respiration rate following IP injection of different pharmaceutical agents. Fish injected with vasotocin had elevated respiration rates compared to saline injected controls, paralleling the pattern seen during recovery of brain injection. * *p* ≤ 0.05; ** *p* ≤ 0.01; *** *p* ≤ 0.001

**Supplemental Figure 4.**
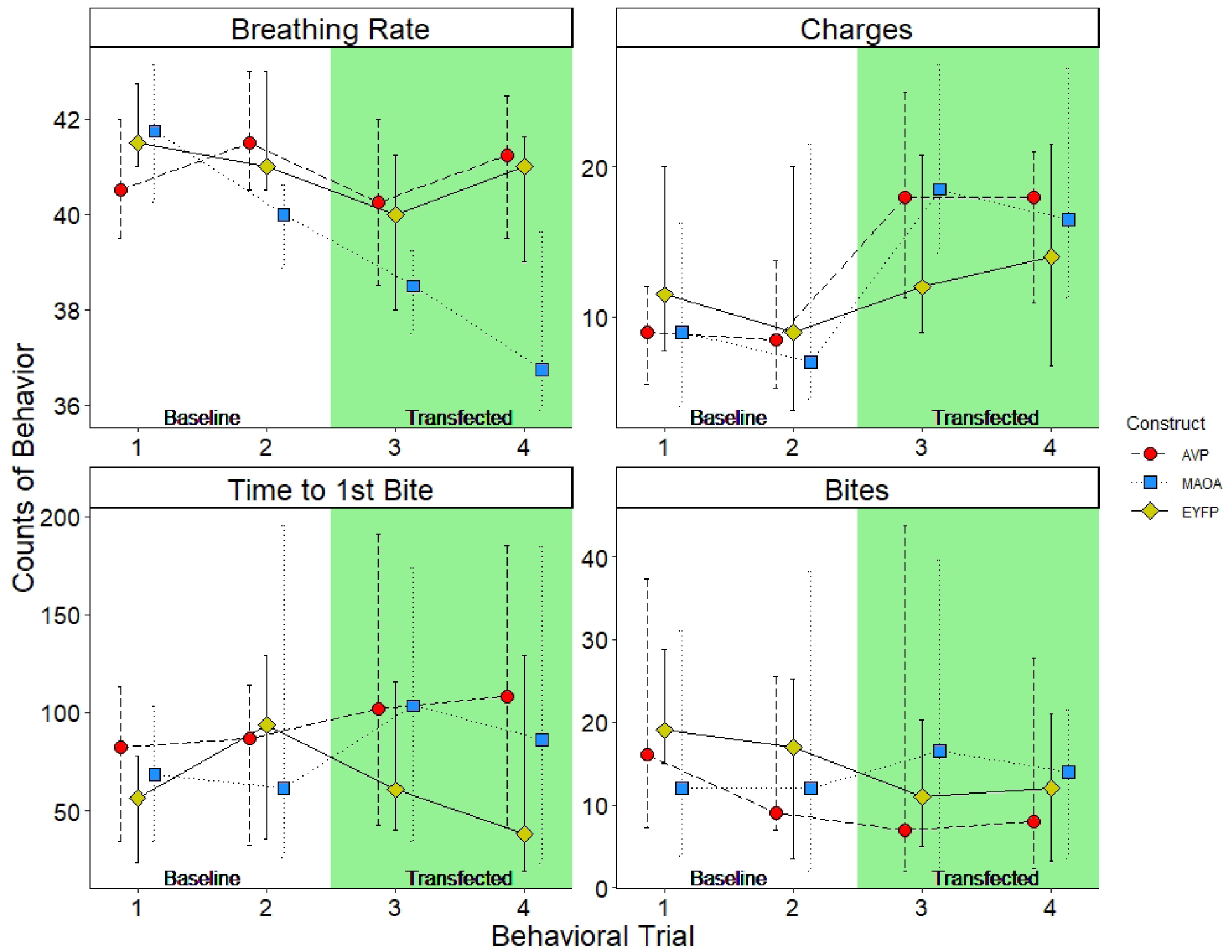
Breathing rate and repeatable behaviors across all trials & constructs. Breathing rate decreased significantly following transfection of only MAOA. Charges increased following AVP or MAOA transfection but not in control EYFP fish. Graph presents medians with interquartile range bars.

**Supplemental Table 1.** Within-subject comparison of territorial aggression following pharmaceutical manipulation of vasotocin signaling compared to baseline. Brain and IP injection results were similar with only the highest dosage altering behavior. No group (including the Manning treatment) significantly differed from saline-injected controls. * *p* ≤ 0.05; ** *p* ≤ 0.01; *** *p* ≤ 0.001

|  | N | Bites |  |  |  | Charges |  |  |  |
| --- | --- | --- | --- | --- | --- | --- | --- | --- | --- |
|  |  | Change | Z | Effect size (r) | P | Change | Z | Effect size (r) | P |
| AVP (Brain, 0.5 $\mu\text{g}/\text{gbw}$) | 3 | -29.3 | -1.3 | 0.93 | 0.18 | -2.0 | -0.9 | 0.82 | 0.35 |
| AVP (IP, 0.5 $\mu\text{g}/\text{gbw}$) | 16 | -10.0 | -1.6 | 0.41 | 0.11 | 1.8 | -1.1 | -0.29 | 0.25 |
| AVP (Brain, 1 $\mu\text{g}/\text{gbw}$) | 2 | -16.5 | -0.9 | 0.95 | 0.37 | -24.5 | -0.9 | 0.95 | 0.37 |
| AVP (IP, 5 $\mu\text{g}/\text{gbw}$) | 5 | -2.4 | -1.4 | 0.66 | 0.18 | -6.3 | -0.8 | 0.43 | 0.41 |
| AVP (Brain, 10 $\mu\text{g}/\text{gbw}$) | 10 | -17.8 | -1.5 | 0.52 | 0.12 | -8.2 | -2.0 | 0.66 | 0.04 * |
| AVP (IP, 10 $\mu\text{g}/\text{gbw}$) | 9 | -2.4 | -0.3 | 0.12 | 0.80 | -6.3 | -2.0 | 0.67 | 0.05 * |
| Manning (IP) | 16 | -32.4 | -2.6 | 0.67 | 0.008 ** | -5.8 | -2.5 | 0.64 | 0.01 * |
| Saline (Brain) | 10 | -8.1 | -1.2 | 0.40 | 0.22 | 1.4 | -0.6 | -0.21 | 0.54 |
| Saline (IP) | 19 | 6.9 | -0.7 | 0.17 | 0.46 | -1.9 | -1.2 | 0.27 | 0.23 |

**Supplemental Table 2.** Repeatability of territorial aggression behaviors and respiration rate across the two trials at baseline and after transfection using two-way mixed, single score ICC (type 3,1) for all fish (*N* = 54) *EYFP* is the control, *AVP* and *MAOA* are genes of interest.

| Measure | Construct | Baseline |  |  | Transfected |  |  | Repeatable at both timepoints |
| --- | --- | --- | --- | --- | --- | --- | --- | --- |
|  |  | ICC | 95% CI | p-value | ICC | 95% CI | p-value |  |
| Respiration Rate | *EYFP* | 0.49 | 0.01, 0.78 | 0.02 | 0.91 | 0.76, 0.97 | 2.51E-07 | Yes |
|  | *AVP* | 0.38 | -0.09, 0.71 | 0.05 | 0.73 | 0.42, 0.89 | 1.76E-04 | Yes |
|  | *MAOA* | 0.53 | 0.12, 0.78 | 0.007 | 0.35 | -0.09, 0.68 | 0.06 |  |
| Time to Orient | *EYFP* | 0 | -0.48, 0.48 | 0.50 | 0.13 | -0.37, 0.58 | 0.30 |  |
|  | *AVP* | 0 | -0.46, 0.46 | 0.50 | 0 | -0.46, 0.46 | 0.50 |  |
|  | *MAOA* | 0.04 | -0.40, 0.47 | 0.43 | 0.21 | -0.24, 0.59 | 0.18 |  |
| Time to 1st Bite | *EYFP* | 0.77 | 0.47, 0.91 | 1.32E-04 | 0.47 | -0.01, 0.78 | 0.03 | Yes |
|  | *AVP* | 0.51 | 0.07, 0.78 | 0.01 | 0.28 | -0.20, 0.65 | 0.12 |  |
|  | *MAOA* | 0.48 | 0.06, 0.75 | 0.01 | 0.74 | 0.45, 0.89 | 6.96E-05 | Yes |
| Bites | *EYFP* | 0.88 | 0.68, 0.95 | 2.0E-06 | 0.78 | 0.47, 0.92 | 1.17E-04 | Yes |
|  | *AVP* | 0.45 | 0, 0.75 | 0.03 | 0.73 | 0.42, 0.89 | 1.69E-04 | Yes |
|  | *MAOA* | 0.62 | 0.25, 0.83 | 0.001 | 0.59 | 0.21, 0.82 | 0.002 | Yes |
| Charges | *EYFP* | 0.64 | 0.23, 0.86 | 0.003 | 0.69 | 0.31, 0.88 | 0.001 | Yes |
|  | *AVP* | 0.55 | 0.12, 0.80 | 0.008 | 0.54 | 0.11, 0.80 | 0.01 | Yes |
|  | *MAOA* | 0.39 | -0.06, 0.70 | 0.04 | 0.83 | 0.62, 0.93 | 1.61E-06 | Yes |
| Trips | *EYFP* | 0.75 | 0.41, 0.90 | 2.82E-04 | 0.32 | -0.19, 0.70 | 0.10 |  |
|  | *AVP* | 0.39 | -0.08, 0.72 | 0.05 | 0.19 | -0.29, 0.59 | 0.22 |  |
|  | *MAOA* | 0.61 | 0.24, 0.82 | 0.002 | 0.66 | 0.31, 0.85 | 6.18E-04 | Yes |

